# Ultrastructural analysis of synapses after induction of spike-timing-dependent plasticity

**DOI:** 10.1101/2025.06.19.660567

**Authors:** Rui Wang, Michaela Schweizer, Margarita Anisimova, Christine E. Gee, Thomas G. Oertner

## Abstract

Repeated sequential activation of connected neurons causes lasting changes in synaptic strength, a process known as spike-timing-dependent plasticity (STDP). Recently, sequential spike patterns have been induced without electrodes, using two spectrally separated channelrhodopsins. However, due to the difficulty of labeling and localizing the few connecting synapses between the stimulated pre and postsynaptic neurons (∼1-5 per neuron pair), ultrastructural analysis after STDP has not been reported. Here, we optogenetically induce STDP at CA3-CA1 hippocampal synapses and identify stimulated boutons and spines in CA1 using transmission electron microscopy (TEM). Presynaptic CA3 neurons express vesicle-targeted horseradish peroxidase, cre recombinase and cre-dependent ChrimsonR, a red light activatable channelrhodopsin. Postsynaptic neurons express violet light activatable CheRiff and dAPEX2, an enhanced ascorbate peroxidase. In transmission electron microscopy, presynaptic boutons and postsynaptic spines are readily identifiable with well-preserved ultrastructural features. Our labeling strategy allows ultrastructural analysis of optogenetically manipulated neurons and their synapses.

## Introduction

Exactly what changes in the brain when a new memory is formed is not fully understood. The memory trace, or “engram”, may consist of minute structural changes at the synaptic connections between neurons that have repeatedly fired together. Spike-timing dependent plasticity (STDP), induced by precisely ordered firing of neurons, causes timing-dependent potentiation (tLTP) or timing-dependent depression (tLTD) of cortical synapses^1–3^. Recently, we have shown that the functional changes induced by optogenetic STDP (oSTDP) in the hippocampus can be very long-lasting^4^. Assessing the strength of synaptic connections days after induction of plasticity became possible by controlling the firing of pre- and postsynaptic action potentials through light-gated channels (channelrhodopsins) instead of invasive recordings with glass electrodes.

It is unknown whether the relative changes in synaptic strength induced by STDP are maintained by physical changes in synapse size. Multiple synapses formed by boutons from one axon onto spines of a specific postsynaptic neuron have remarkably similar ultrastructure, suggesting that synaptic structure is tightly controlled by the activity history in the presynaptic and postsynaptic neurons^5–7^. However, these studies did not examine the actual activity history or strength of the connection between the pre and postsynaptic neurons. An increase in synaptic strength associated with tLTP could equally be maintained by either an increase in size of existing synapses, or by forming additional synaptic connections between neurons. The latter is supported by the observation that growth of new spines can be induced by localized glutamate release onto a dendrite^8,9^.

Since the size of a typical CNS synapse is close to the diffraction limit of light, subcellular synaptic structures cannot be visualized by conventional light microscopy. Stimulated emission depletion (STED) microscopy is a promising approach to image dendritic spines and associated presynaptic vesicle clusters^10^. However, fluorescence-based methods alone visualize only labeled entities and provide no information about surrounding unlabeled structures or cells in the tissue, unless they are combined with electron microscopy (EM)^11^. After osmium tetroxide staining, unlabeled membranes become visible in EM, providing precise information about the volume of spines and presynaptic boutons, the size of postsynaptic densities and active zones, and the presence of organelles such as vesicles, mitochondria and endoplasmic reticulum. Several studies have analyzed synaptic ultrastructure after high frequency electrical stimulation^12 13 14 43^. A limitation of blind electrical stimulation is that just a subset of axons is activated at the desired frequency, leading to a mix of potentiated, partially stimulated and unstimulated synapses in the analyzed samples.

Correlative light-electron microscopy (CLEM) allows analyzing the ultrastructure of identified synapses that were individually activated, e.g. by glutamate uncaging in Mg^2+^-free ACSF^15^. The resulting rapid changes in postsynaptic ultrastructure are impressive, but it is unclear whether similar changes can be induced by physiological activity of individual presynaptic axons. Optogenetic stimulation offers the exciting possibility to drive spiking in identified neurons non-invasively under physiological conditions. It has recently been combined with membrane-targeted peroxidase expression to identify light-activated axons by post-embedding immunogold labeling^16^. Our goal was to extend this promising approach to establish a labeling protocol that would allow the detection of STDP-stimulated (“paired”) synapses and the quantification of activity-related ultrastructural changes. Paired synapses could then be compared to three types of “control” synapses within the same section: 1) nearby synapses between unlabeled/unpaired neurons, 2) synapses from the same presynaptic neuron onto unstimulated postsynaptic neurons, 3) other synapses onto the same postsynaptic dendrite. Control synapses within the same tissue block are ideal to exclude small variations in fixation or staining that could affect the quantification of synaptic ultrastructure. We present a methodological workflow starting with viral transduction of neurons, induction of optogenetic STDP, and processing of fixed tissue for transmission electron microscopy (TEM) to study ultrastructural changes at synapses associated with tLTP.

Novel is the combination of opsins, EM compatible labels and staining procedure that allows unambiguous identification of presynaptic boutons of ChrimsonR-expressing CA3 neurons and postsynaptic spines and dendrites of CheRiff-expressing CA1 neurons. Since both EM labels are peroxidases that are detected by DAB staining, the combination of labels we present here is also compatible with serial block-face imaging, automated 3D reconstruction and analysis^17–20^, which should greatly improve the throughput and success rate of finding potentiated synapses. Since both labels are genetically expressed, they can be precisely targeted to the cell/types of interest and are broadly applicable to studies of other synapses of interest. Postsynaptic dAPEX2 could also be expressed in other cell types, such as astrocytes or microglia, if the goal is to study their proximity to specific synapses. There is no reliance on antibody-mediated labeling, so there are no concerns about tissue penetration, antigen availability, selectivity or specificity. In addition, fluorescence does not need to be retained during fixation steps, so these can be strong and optimized for easy identification of membranes and organelles. We expect this method to be particularly useful in answering open questions about the fate of synapses and their ultrastructural changes after STDP.

## Results

### The ultrastructure of potentiated synapses

Our goal was to induce spike-timing-dependent plasticity (STDP) non-invasively and subsequently examine the ultrastructure of the presynaptic boutons and postsynaptic spines connecting the co-stimulated neurons (Fig. 1). The most critical step in the development of this workflow was the identification of a pair of peroxidase-based genetic tags that, after processing and transmission electron microscopy (TEM), would allow the unambiguous identification of the structures originating from presynaptic and postsynaptic neuron. We focused on Schaffer collateral synapses formed by axons of CA3 pyramidal cells onto spines of CA1 pyramidal cells in rat organotypic hippocampal slice cultures. Since the slice cultures are ∼300 µm thick, we wanted to avoid antibody-based labeling, which often fails to reveal structures more than ∼50 µm below the surface.

**Figure 1:**
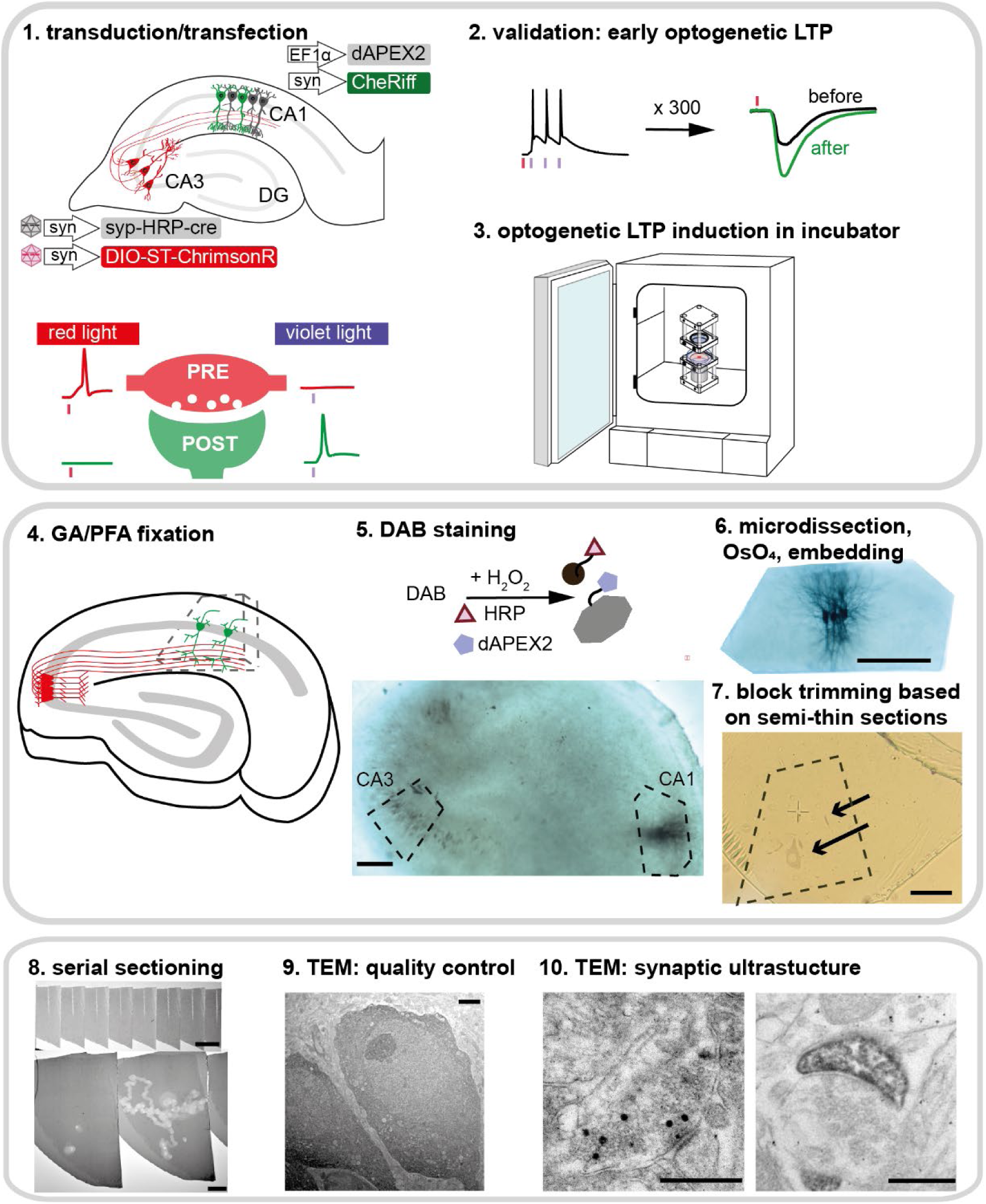
Workflow of STDP induction and sample preparation to study synaptic ultrastructure. **Step 1**: AAV-assisted transduction of presynaptic neurons and single cell electroporation to transfect postsynaptic neurons. Red light (635 nm) only induces action potentials in presynaptic neurons expressing ChrimsonR, violet light (405 nm) only induces action potentials in postsynaptic neurons expressing CheRiff. **Step 2 and 3**: Optogenetic induction of STDP. Induction of all-optical tLTP by 300 repetitions of single presynaptic (red) and three postsynaptic (violet) action potentials. Validation of all-optical tLTP at early and late timepoints. **Step 4**: Fixation at the desired time point after induction of tLTP. **Step 5**: Metal-enhance DAB staining to reveal HRP in presynaptic vesicles and dAPEX2 in the postsynaptic neurons. **Step 6**: Micro-dissection of the stained CA3 and CA1 regions, osmification and embedding. **Step 7**: Further trimming of the block based on DAB-stained somata and processes in semi-thin sections (arrows). **Step 8**: Ultra-thin serial sectioning for TEM. **Step 9**: Verify structural preservation/sample quality at the soma area of labeled CA3 and CA1 neurons. **Step 10:** Locating and imaging structures-of-interest. Scale bars: 5 and 6: 200 μm; 7:20 μm; 8: 60 μm, 20 μm; 9: 2 μm; 10: 500 nm.

### Step 1: Viral transduction of CA3 neurons and single-cell electroporation to transfect CA1 neurons

To independently spike and label presynaptic CA3 neurons, we used two recombinant adeno-associated viral vectors (rAAVs). One rAAV encoded the red-light activatable channelrhodopsin ChrimsonR, the second a synaptic vesicle-targeted horseradish peroxidase (synaptophysin-HRP, syp-HRP). Initial attempts to package a single plasmid encoding ChrimsonR-tdTomato and syp-HRP into one rAAV were unsuccessful. When we mixed two rAAVs with capsid serotype rh10, the percentage of neurons expressing both proteins was disappointingly low (∼40%). We solved this problem of poor co-expression with the Cre/LoxP system^21^. We packaged a cre-dependent version of ChrimsonR in one rAAV and a bicistronic vector for co-expression of synaptophysin-HRP and cre (syp-HRP-IRES-cre) in a second one, both with capsid serotype AAV9. We observed that one week after local injection of the mixed rAAVs, 96% of transduced CA3 neurons expressed both ChrimsonR and syp-HRP-IRES-cre (Fig. 2a, b). The remainder (4%) appeared to express only ChrimsonR, indicating that cre expression was below the antibody detection limit. This interpretation is supported by the observation that no neurons expressed ChrimsonR when the rAAV was applied alone (Fig. 2c). Somewhat surprisingly, we detected no neurons expressing only syp-HRP-IRES-cre when the two rAAVs were applied together. Taken together, we expect almost no cases where light-activated CA3 neurons do not express the EM label syp-HRP and vice versa. A small volume of the virus mix was pressure-injected into area CA3 of hippocampal cultures using a picospritzer (1.8 bar, 50 ms pulse). As we have previously shown^4^, it is very important that the number of transduced CA3 neurons is in the range of 30-40 to achieve successful and specific induction of oSTDP. Simultaneous activation of a larger number of CA3 neurons causes firing of secondary (non-transfected) neurons and in consequence, synaptic plasticity at unlabeled synapses. For reliable oSTDP, light-induced EPSCs in the postsynaptic neuron should be well below 200 pA (Fig. S2).

**Figure 2:**
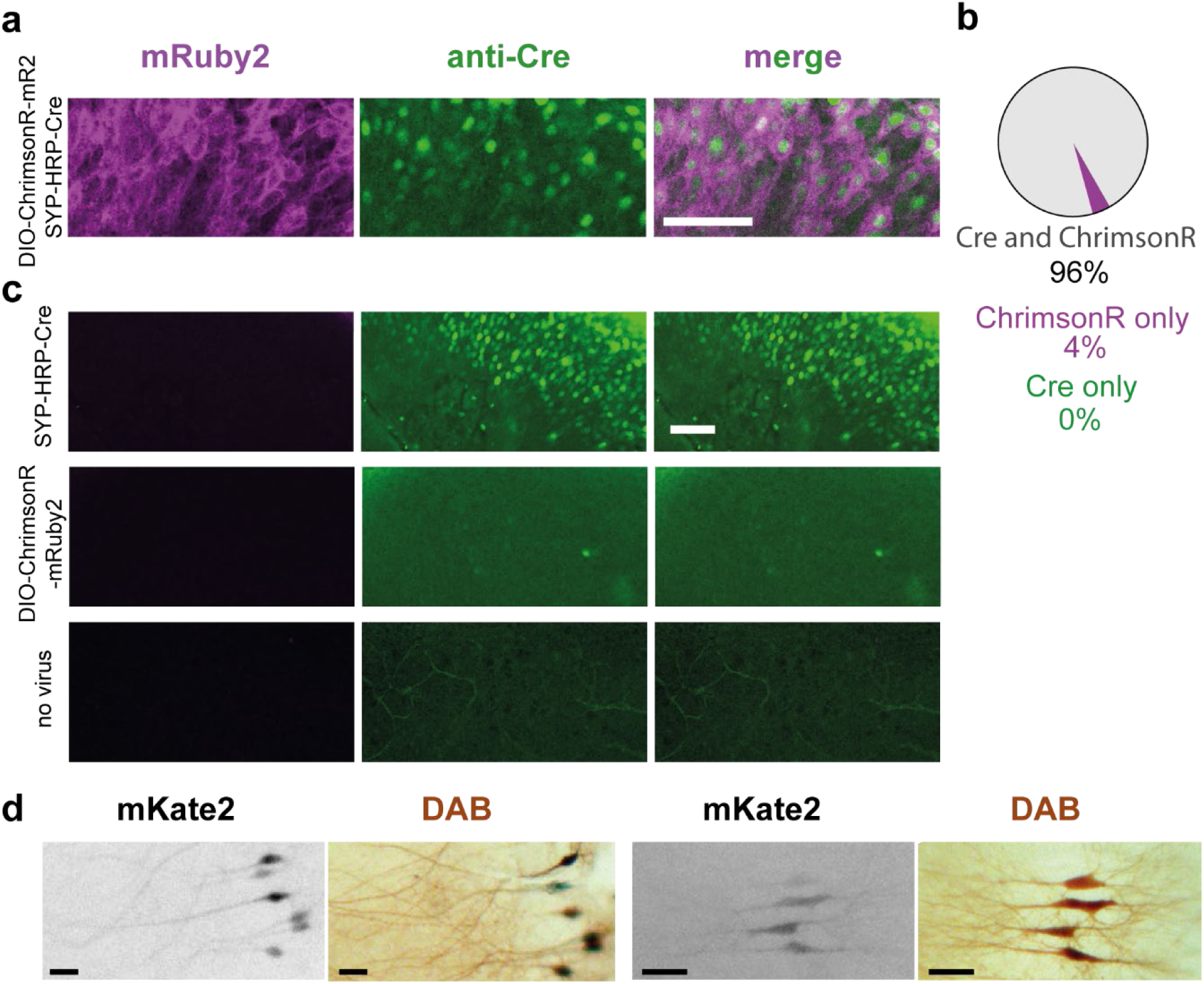
Assessment of electron microscopy label and channelrhodopsin co-expression. (**a**) CA3: A mixture of rAAV^2/9^-syn-DIO-ChrimsonR-mRuby2-ST and rAAV^2/9^-syn-SYP-HRP-IRES-Cre was co-injected into CA3. Slices were fixed and immunostained against mRuby2 (magenta) and cre (green). Scale bar: 100 μm. (**b**) Quantification of co-expression rate of ChrimsonR-mRuby2 and syp-HRP-IRES-Cre. N = 202 cells (4 slice cultures). (**c**) Application of the individual viruses or no virus. Scale bar: 100 μm. (**d**) Two examples of co-expression in CA1 pyramidal neurons electroporated with plasmids encoding dAPEX2, mKate2 and CheRiff-eGFP: mKate2 fluorescence (contrast-inverted grayscale) several days before fixation corresponded to the neurons revealed by DAB staining (brown). Scale bars: 40 μm (left images), 70 μm (right images).

To independently spike and label a few postsynaptic CA1 neurons, we co-expressed the violet light activatable channelrhodopsin CheRiff with dAPEX2^20^, an enhanced version of soluble APEX2^22^, by single-cell electroporation of a 1:20 plasmid mixture^23^. A high concentration of dAPEX2 was critical to later trim the resin block close to the dark dendritic branches of electroporated cells. For electrophysiology experiments, a small amount of mKate2 plasmid was added to allow us to visualize the transfected CA1 neurons without activating CheRiff. In our patch-clamp experiments, we never encountered mKate2-expressing neurons without photocurrents, suggesting highly reliable co-expression of the electroporated plasmids. Importantly, neurons with even rather dim mKate2 fluorescence were revealed by DAB staining (Fig. 2d).

### Step 2: Validation of all-optical tLTP induction

For optogenetic STDP induction, we used two channelrhodopsins with largely separated activation spectra, ChrimsonR and CheRiff ^24,25^. Previously, we carefully characterized their properties and found that low intensity violet light pulses (∼1 mW mm^−2^, 2 ms) selectively generate single action potentials in CheRiff-expressing neurons while high intensity orange to red light pulses (∼7 mW mm^−2^, 2 ms) selectively spike ChrimsonR-expressing neurons^4^ (Fig. 1, step 1). Slice cultures with virally transduced CA3 neurons and electroporated CA1 neurons (Fig. 3a) were stimulated with a 594 nm laser positioned though the condenser to illuminate CA3, or a red LED coupled through the objective (Fig. S1). CheRiff-expressing CA1 neurons were stimulated by a 405 nm LED coupled through the objective. Using the exact combinations of viral vectors and plasmids described in Step 1, we verified that ChrimsonR/syp-HRP expressing CA3 neurons faithfully followed red light flashes with single action potentials in cell-attached recordings (Fig. 3b). When recording from CA1 neurons, a 2 ms, 594 nm light pulse elicited an excitatory postsynaptic current (EPSC) of 100-200 pA. To induce timing-dependent LTP (tLTP), the 594 nm light pulse was paired with a burst of 3 violet light flashes, delayed so that the first spike in the CA1 neuron occurred 10 ms after the onset of the EPSC (+10 ms causal pairing). Repeated 300 times at 5 Hz, this spike/burst stimulation pattern produced EPSCs and 3 spikes in CheRiff/dAPEX2-expressing CA1 neurons and induced tLTP (Fig. 3c-d).

**Figure 3:**
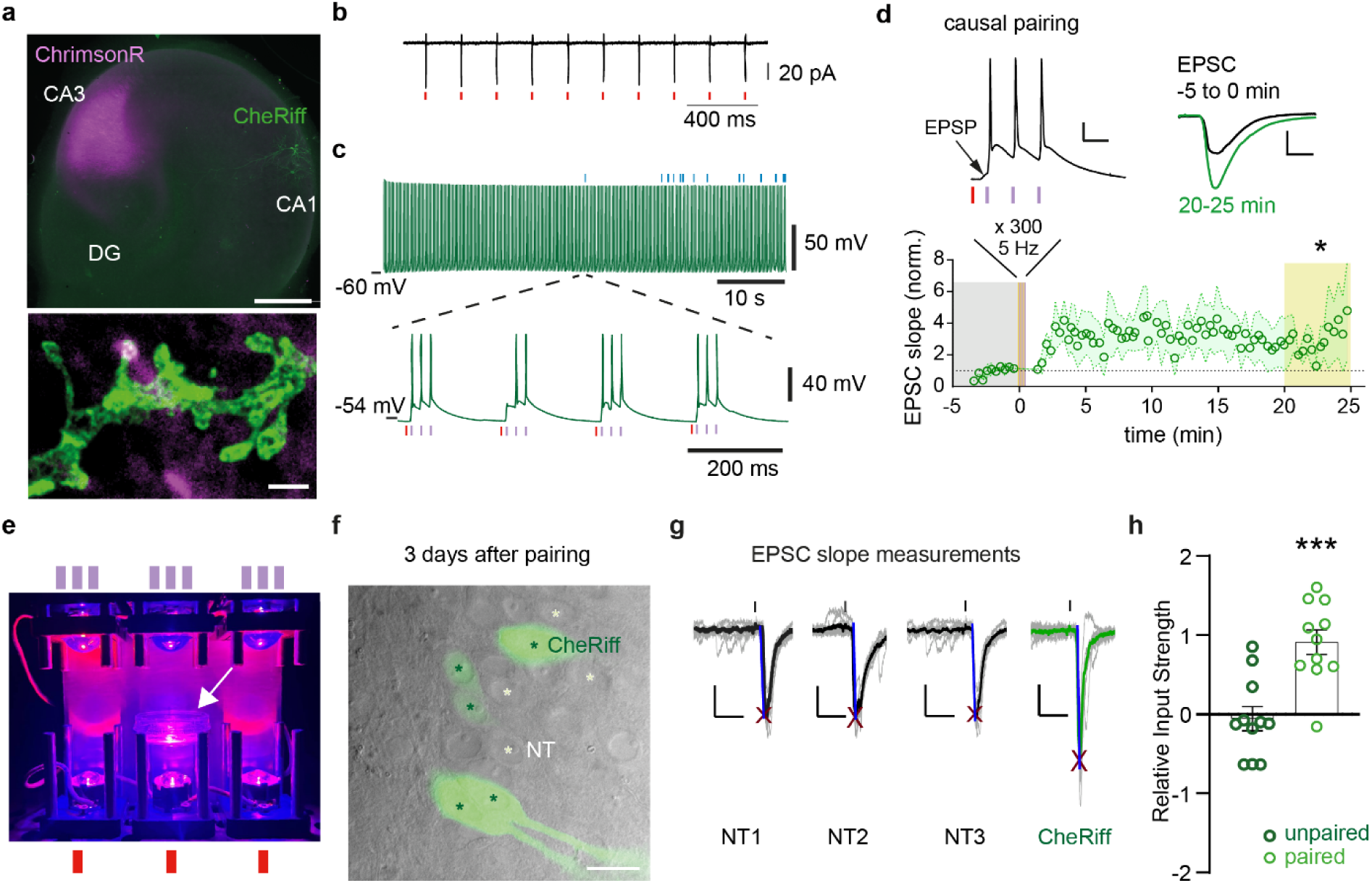
All-optical tLTP in hippocampal slice culture lasts for 3 days. (**a**) Transfection/transduction strategy for hippocampal STDP. Top: Wide-field image of hippocampal slice culture shows cluster of virally transduced CA3 pyramidal cells (magenta) and single transfected CA1 pyramidal cell (green). Bottom: STED image (z projection) of CheRiff-expressing dendrites (green) overlaid with confocal image of presynaptic ChrimsonR-expressing boutons (magenta), showing a putative synaptic contact (white). Scale bars: 500 μm, 1 μm. (**b**) Representative cell-attached recordings of Flex-ST-ChrimsonR, co-expressed with SYP-HRP-Cre CA3 neurons, recorded in culture medium. CA3 Syp-HRP-Cre + Flex-ST-ChrimsonR neuron fires single action potential is response to red light pulses (red tics). (**c**) Representative current-clamp recording of a CA1 neuron while applying paired optical stimulation. Blue ticks above the trace indicate stimulations where only 2 postsynaptic spikes were triggered instead of 3 (see expanded timescale below). In this example, 286 out of 300 paired stimulations were flawless. (**d**) After oSTDP induction (left insert, scale bars: 20 mV, 20 ms), the slope of EPSCs increases (right insert, scale bars: 200 pA, 10 ms). Red arrow indicates the light pulse (7 ms conduction delay from CA3 to CA1). Paired t-test, P = 0.044, n = 4 slice cultures. Data plotted as mean ± SEM. (e) Three LED towers for in-incubator stimulation. The middle tower contains a 35 mm petri dish with cell culture insert (white arrow). Ticks symbolize the three 405 nm light pulses from the top LED and single 635 nm pulse from the bottom LED. (**f**) Dodt contrast and epifluorescence image of CA1. CheRiff neurons (green asterisks) were distinguished from non-transfected (NT) neighbors (white asterisks) by their somatic fluorescence (green). Scale bar: 36 μm. (**g**) Calculation of input strength. Vertical lines indicate presynaptic optical stimulation (1 ms, 594 nm, identical light intensity was used to stimulate sequentially patched NT and CheRiff cells in each slice). For each neuron, the 20% to 60% slope was measured on the averaged EPSC (thick traces) from 10 consecutive stimulations (gray traces). Red crosses show auto-detected peaks. Scale bars: 25ms, 50 pA. (**h**) Effect of paired optical stimulation on input strength. Each circle shows the EPSC slope of a CheRiff-expressing neuron normalized to EPSC slope in 2-3 non-transfected neighbors. The tLTP group (light green) was assessed 3 days after causal pairing, the unpaired group (dark green) received no paired stimulation. Unpaired t-test, P = 0.0002, n = 11, 11 neurons. Bars show mean ± SEM.

### Step 3: Optogenetic stimulation in the incubator

To follow the stimulated synapses over several days, or before fixing for ultrastructural analysis, we optically stimulated slice cultures inside the cell culture incubator (Fig. 3e). As there is no baseline to compare EPSCs after STDP, the EPSC slope of paired CA1 neurons is normalized to the average EPSC slope of several (3-5) neighboring non-transfected neurons (NT EPSC, Fig. 3f, g). The relative input strength is calculated as:

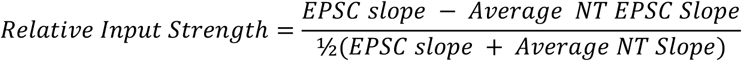

As previously reported^4^, the relative synaptic input strength to CheRiff-expressing CA1 neurons was stronger compared to nearby non-transfected neurons 3 days after causal pairing (Fig. 3g, h). Thus, in organotypic slices, tLTP persists at least 3 days (late tLTP). Importantly, too many simultaneously spiking CA3 neurons will directly drive spiking in the postsynaptic CA1 neurons, even in the absence of CheRiff and violet light^4^. In this case, the NT neurons will also undergo STDP, which means the specificity that only connections between the ChrimsonR-expressing presynaptic neurons and the CheRiff-expressing postsynaptic neurons are potentiated will be lost. LTP is still induced under these conditions, but it is not possible to use unlabeled postsynaptic neurons as a reference group. Thus, if the goal is to compare the ultrastructure of “paired” synapses with that of neighboring “control” synapses in CA1, it is a requirement that relatively few CA3 neurons express ChrimsonR/syp-HRP.

### Steps 4-9: Preparation of tissue for TEM and serial sectioning

Ten minutes after plasticity induction, each slice culture was fixed and processed for EM examination. We tested several staining protocols for syp-HRP and dAPEX2. We found that metal-enhanced DAB staining followed by osmification was necessary to reliably visualize both the axonal boutons of CA3 neurons and the spines and dendrites of CA1 neurons. Briefly, after the DAB reaction using metal-enhanced DAB substrate (Thermo 34065), the slice was trimmed under visual control using the stained CA1 neurons as a guide. The trimmed specimens were then fixed with 1% (w/v) osmium tetroxide, dehydrated in an ascending series of ethanol, and embedded in Epon 812. Several rounds of trimming were required to center the brown DAB-stained CA1 neurons in the blocks, minimizing the area to be searched for synaptic contacts. Semi-thin sections were cut and immediately inspected for DAB-stained somata (Fig. 1, Step 7). After further trimming of the edges of the mounted block, serial ultrathin sections were cut. For quality control, we sectioned separate blocks containing the somata of stained CA3 and CA1 neurons and inspected them for abnormalities (Figs. S3, S4). Organelles such as Golgi apparatus, mitochondria and rough ER were abundant in the cytoplasm and nuclei appeared normal. Note that dAPEX2 creates a granular precipitate of electron-dense particles in the nuclei and cytoplasm of CA1 neurons that is not associated with any particular biological structure (Fig. S4).

### Step 10: Analysis of synaptic ultrastructure after STDP

Labeled synaptic vesicles in channelrhodopsin-expressing axons appeared as distinct electron-dense dots. This allowed for the identification of stimulated axons and their boutons (Fig. 4). The somata (Fig. S4), dendrites and spines of HRP-labeled postsynaptic neurons were filled with granular electron-dense material that was easy to recognize in ultrathin sections. Potential synaptic connections were identified as contacts between labeled presynaptic and postsynaptic compartments (Fig. 4a, Fig. S5). Both single- and double-labeled synapses could be identified in the same specimen (Fig. 4b), which provided important internal controls: Contacts of labeled boutons with unlabeled spines experienced presynaptic activity and glutamate release, but not paired with postsynaptic spikes. Labeled spines in contact with unlabeled boutons experienced triplets of back-propagating action potentials, but not paired with any presynaptic activity. These forms of activity are not expected to induce strong long-term plasticity, though some degree of heterosynaptic plasticity is possible. As a second oSTDP group, we analyzed synapses from a slice culture that we stimulated in anti-causal sequence: Three postsynaptic action potentials were followed by one presynaptic action potential (Fig. S6). Anti-causal stimulation is known to induce early timing-dependent LTD (tLTD)^4^.

**Figure 4:**
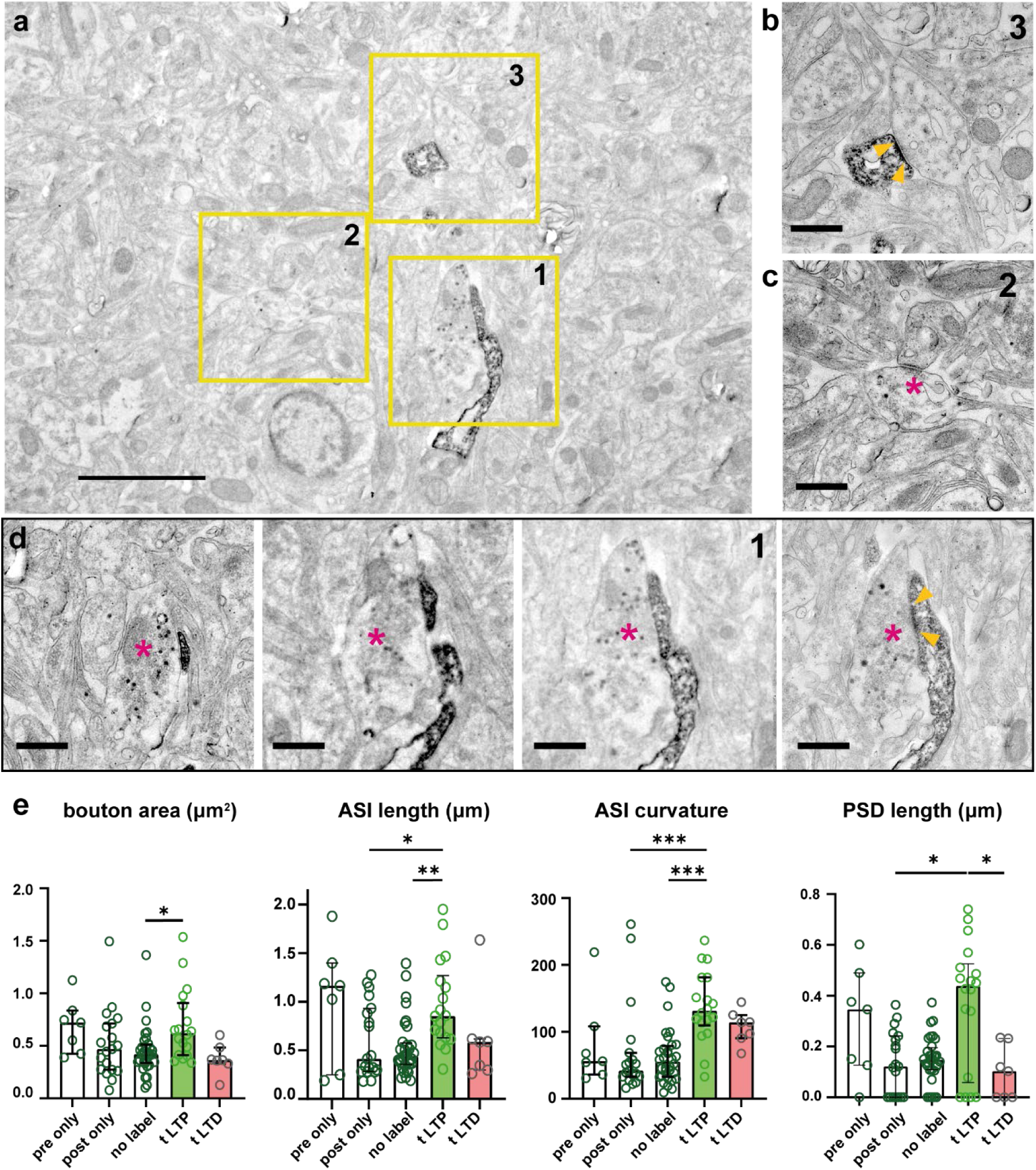
Causal pairing drives expansion of spine-bouton interface and postsynaptic density. (**a**) In the same FOV, a double labeled synapse (1) and single labeled synapses (2, 3) are visible. (**b**) High resolution image of the labeled spine (3) in synaptic contact with unlabeled bouton. Yellow arrow tips delimit postsynaptic density (PSD). (**c**) High resolution image of bouton (2, magenta asterisk) with two labeled vesicles in contact with unlabeled postsynaptic element. (**d**) Serial sections through the double labeled synapse (red asterisk: bouton, blue arrow: spine). The postsynaptic density is visible in the last section (yellow arrow heads). Scale bars: 2 μm, 500 nm. (**e**) Analysis of bouton area, axon-spine interface (ASI) length, ASI curvature, and PSD length (pre only: n = 7; post only: n = 19; no label: n = 32; tLTP: n = 18; tLTD: n = 7 synapses). Bars show median ± quartiles. ANOVA followed by Kruskal Wallis tests (*p < 0.05, **p < 0.01, ***p < 0.001).

A comparison of the size of presynaptic boutons revealed that tLTP synapses had a larger cross-sectional area than unlabeled synapses (Fig. 4e), resulting in longer axon-spine interfaces (ASI). Interestingly, the ASI became strongly curved after tLTP, suggesting that the activated spine heads began wrapping around the active presynaptic boutons (Fig. 4e). Actin polymerization inside spine heads is indeed triggered by NMDAR-mediated calcium influx^26^ and enlarged, cup-shaped spines have frequently been reported after LTP protocols^27,28^. Not all bouton-spine contacts had a clearly identifiable postsynaptic density (PSD) in the inspected sections (Fig. S5). However, the median length of double-labeled PSDs after tLTP increased significantly compared to PSDs in contact with non-activated boutons (post only, Fig. 4e). The PSDs of tLTP synapses were also larger than the PSDs of the tLTD group, reflecting the well-established functional consequences of causal versus anti-causal stimulation^1,4^. Due to our limited 2D dataset, we could not analyze spine morphology in more detail. Since LTP is believed to affect the geometry of the spine neck^29^, generating 3D data of synapses after oSTDP is an exciting prospect.

In summary, the task of finding the connecting synapses between a small number of presynaptic neurons and a specific CA1 neuron is difficult, but not impossible. Based on the EPSP amplitude in response to presynaptic optogenetic stimulation, a typical mEPSC amplitude of 20 pA^30^ and an average release probability of 0.5, we estimate ∼20 light-stimulated synapses onto a single postsynaptic neuron, corresponding to about 0.1-0.2 % of spines^31^. Thus, approximately 1000 spines from a single CA1 neuron needed to be inspected to find one double-labeled synapse. We focused our search on *stratum radiatum*, where CA3 axons form most of their contacts onto the oblique dendrites of CA1 pyramidal neurons.

## Discussion

We present a workflow to first induce timing-dependent LTP with light pulses and subsequently localize and study potentiated Schaffer collateral synapses in ultrathin seral sections. Compared to previous approaches to study the ultrastructure of potentiated synapses, our method has several advantages: oSTDP is induced inside a cell culture incubator at 37°C, avoiding the insertion of electrodes and other perturbations between plasticity induction and fixation. The resulting tLTP lasts for days, providing a remarkably long time window for studying ultrastructural changes. Using targeted peroxidase labels, each labeled synaptic structure could be assigned unambiguously to presynaptic or postsynaptic neurons and the associated stimulation pattern. This made it easy to identify essential control groups of synapses (presynaptic stimulation only, postsynaptic stimulation only, unstimulated) present within each tissue block. DAB staining and sample trimming under a bright-field microscope are basic procedures, correlative fluorescence imaging or antibody staining was not required for our approach. Thus, we could use a strong fixation protocol to obtain high contrast images. All viral vectors are compatible with the lowest biosafety level (BSL-1). By combining optogenetic tools with peroxidase-based genetic tags using Cre/LoxP, we present a method to study the ultrastructural changes of tLTP from minutes to days after plasticity induction. As a future perspective, it may be possible to induce oSTDP in a suitably equipped high-pressure freezing machine^32^ to study rapid changes in synaptic ultrastructure.

### Limitations of the study

The main limitation of our method is its low throughput, limited by the time required to manually search and locate double-labeled synapses with conventional serial section TEM. Deep learning-based methods have proven extremely powerful for the automated tracing and detection of synapses in EM images^33,34^ and could be applied to this problem. A further advance would be to combine our stimulation and labeling approach with volume EM methods such as array electron tomography^35^, serial block face EM^36^ or focused ion beam milling^37,38^. Since DAB staining of tissue blocks is straightforward (no antibody staining required), our approach seems particularly well suited for these high throughput methods. We did not, however, test this idea. A second limitation is the possibility of missing small presynaptic boutons of labeled CA3 neurons, as only about 20 % of presynaptic vesicles become electron-dense^20^. The comparatively large size of labeled boutons (pre-only; Fig. 4e) is probably a reflection of this bias. Tracking axons through serial sections or 3D volumes will mitigate this risk, as nearby boutons containing labeled vesicles will signal that an adjacent unlabeled bouton was also active during plasticity induction.

## Resource availability

### Lead contact

Requests for resources, reagents and further information should be directed to the lead contact, Thomas G. Oertner.

### Materials availability

The pAAV-Syn-SYP-HRP-IRES-Cre plasmid generated in this study has been deposited to Addgene (#231252). Recombinant AAVs used in this study are available from the lead contact upon request.

### Data and code availability

- TEM images and electrophysiological data generated during this study will be shared by the lead contact upon request.
- Custom code for electrophysiology analysis has been deposited at GitHub and is publicly available as of the date of publication (see key resources table).
- Any additional information required to reanalyze the data reported in this paper is available from the lead contact upon request.

## Acknowledgments

We thank Jan Schröder, Iris Ohmert, Emanuela Szpotowicz and Saskia Siegel for their excellent technical assistance. Bas von Bommel performed STED microscopy at the UKE Microscopy Imaging Facility with Virgilio Failla. Ingke Braren from the UKE Vector Facility produced rAAVs. pAAV-hSyn-DIO-ChrimsonR-mRuby2-ST was a gift from Hillel Adesnik (Addgene plasmid # 105448). pAAV-syn-SYP-HRP (Addgene plasmid # 117185) and pAAV-EF1α-dAPEX2 (Addgene plasmid #117173) were gifts from David Ginty. CheRiff (Addgene #51697) was a gift from Adam Cohen, ChrimsonR (Addgene #59171) was a gift from Edward Boyden. Funding was received from the Deutsche Forschungsgemeinschaft (DFG) – project numbers 538090107; 278170285 (FOR 2419); 404539526 (SFB 1328); 500012246 (SPP 2395) – and by the European Research Council (ERC-Synergy 951515).

## Author contributions

TGO, MS, and CEG conceived the study, obtained funding, directed the work and wrote the first draft together with RW. RW performed the experiments, tested plasmids and viruses, prepared tissue including DAB staining and trimming for EM, analyzed data and produced the figures. MS produced serial sections and performed EM, MA established oSTDP. All authors edited and approved the final manuscript.

## STAR Methods

### Key resources table

| REAGENT or RESOURCE | SOURCE | IDENTIFIER |
| --- | --- | --- |
| **Antibodies** |  |  |
| ATTO 647N conjugated (camelid) anti-GFP | NanoTag | Cat# N0304-At647N-S, RRID: AB_2905516 |
| Abberior STAR 580 (camelid) anti-RFP | NanoTag | Cat# N0401-Ab580-S, RRID: AB_2744601 |
| Anti-Cre (mouse) | Millipore | Cat# MAB3120, RRID: AB_2085748 |
| Anti-mouse Alexa Fluor 647 (goat) | Invitrogen | Cat# A-21236, RRID: AB_2535805 |
| **rAAVs** |  |  |
| AAV<sup>2/9</sup> Syn-SYP-HRP-IRES-Cre | UKE | n.a. |
| AAV<sup>2/9</sup> hSyn-DIO-ChrimsonR-mRuby2-ST | UKE | n.a. |
| AAV<sup>2/Rh10</sup> Syn-ChrimsonR-tdT | UKE | n.a. |
| **Chemicals, peptides, and recombinant proteins** |  |  |
| Triton-X-100 | Sigma-Aldrich | X100 |
| PFA | Laborimpex | E15710 |
| GA | Laborimpex | E16320 |
| Osmium Tetroxide | Laborimpex | E19110 |
| Sodium Cacodylate | Scientific Laboratory Supplies | C4945 |
| Goat serum | Capricorn Scientific | GOA-1A |
| Glycidyl ether | Carl Roth | 8619.2 |
| MNA | Carl Roth | 8639.2 |
| DBA | Carl Roth | 8623.2 |
| DMP-30 | Carl Roth | 8621.1 |
| **Critical commercial assays** |  |  |
| Metal Reinforced DAB Substrate Kit | Thermo Fisher | Cat# 34065 |
| **Experimental models: Organisms/strains** |  |  |
| Wild-type Wistar (HsdCpb:Wu,Envigo) or Sprague-Dawley (Hsd:Sprague Dawley®SD® Inotiv) rats | Envigo | HsdCpb: Wu |
| **Recombinant DNA** |  |  |
| pAAV-hSyn-CheRiff-eGFP | Addgene | Cat# 51697 |
| pAAV-Syn-ChrimsonR-tdT | Addgene | Cat# 59171 |
| pCl-Syn-mKate2-N | this study | Modified from Evrogen plasmid Cat# FP182 |
| pAAV-Syn-SYP-HRP-IRES-Cre | this study, Addgene | Cat #231252 |
| pAAV-hSyn-DIO-ChrimsonR-mRuby2-ST | Addgene | Cat#105448 |
| pAAV-EF1α- dAPEX2 | Addgene | Cat#117173 |
| **Software and algorithms** |  |  |
| MATLAB | MathWorks | https://www.mathworks.com |
| Ephus | Suter et al<sup>39</sup> | http://www.ephus.org |
| GraphPad Prism 10 | Dotmatics | https://www.graphpad.com |
| Adobe Illustrator | Adobe | https://www.adobe.com |
| Imaris | OXFORD Instr. | https://imaris.oxinst.com |
| Fiji ImageJ | NIH | https://fiji.sc |
| Electrophysiology analysis tools | This lab | https://github.com/ruiwang1127/Ephys.git |
| Kappa | Gary J. Brouhard lab<sup>40</sup> | https://github.com/fiji/Kappa |

### Experimental model and study participant details

#### Animals

Wild-type Wistar or Sprague-Dawley rats were housed and bred at the University Medical Center Hamburg-Eppendorf (UKE) animal facility and sacrificed according to German Law (Tierschutzgesetz der Bundesrepublik Deutschland, TierSchG), with approval from the Behörde für Justiz und Verbraucherschutz (BJV)-Lebensmittelsicherheit und Veterinärwesen Hamburg and the animal care committee of the UKE.

### Method details

#### Organotypic slice culture

Hippocampus slice cultures were prepared and kept in an antibiotic-free medium as previously described ^41^. Neonatal rat pups at age P4 to P6 under 80 % CO_2_/20 % O_2_ anesthesia were decapitated, and hippocampi were dissected, leaving a portion of the entorhinal cortex attached, in ice-cold sterile slice culture dissection medium (in mM: 248 sucrose, 26 NaHCO_3_, 10 glucose, 4 KCl, 5 MgCl_2_, 2 kynurenic acid, and 0.001% phenol red. pH∼7.4, osmolality 310 to 320 mOsm/kg, bubbled with sterile filtered 95% O_2_/5% CO_2_). 400 μm thick slices were cut from the dissected tissue with a McIllwain tissue chopper (Stoelting). Slices were cultured on 30 mm diameter porous membranes (Millipore PICMORG50), supplied with 1 mL of culture medium (for 500 mL: 394 mL Minimal Essential Medium (MEM, Sigma M7278), 100 mL heat-inactivated donor horse serum (Sigma H1138), 1 mM L-glutamine (Gibco 25030-024), 0.01 mg/mL insulin (Sigma I6634), 1.45 mL, 109 mM NaCl (Sigma S5150), 2 mM MgSO_4_ (Fluka 63126), 1.44 mM CaCl_2_ (Fluka 21114), 6 µM ascorbic acid (Fluka 11140), and 13 mM D-glucose (Fluka 49152), sterile filtered) in a cell culture incubator (37 °C, 5% CO_2_).

#### Transduction of neurons in rat hippocampal slice culture

Transfection was performed at 10-14 days in vitro (DIV) by CA1 single cell electroporation (pAAV-hsyn-CheRiff-eGFP 0.5 ng/μL, pCI-syn-mKate2 10 ng/μL, and dAPEX2 10 ng/μL) and transduction by CA3 local AAV injection (AAV^2/9^-hSyn-DIO-ChrimsonR-mRuby2-ST, 1.05 × 10^14^ vg/mL and AAV^2/9^-SynSYP-HRP-IRES-Cre, 6× 10^13^vg/mL, measured by the WPRE sequence) according to published procedures^23,42^. In Figure 3g, h, CA3 neurons were transduced with AAV^2/Rh10^-syn-ChrimsonR-tdTomato 4.38 × 10^13^ vg/mL and CA1 with hsyn-CheRiff-eGFP 0.5 ng/μL, pCI-syn-mKate2 10 ng/μL. Slice culture medium was partially changed (∼2/3) every 2 to 3 days under dim yellow light (Osram LUMILUX CHIP control T8) to avoid unwanted activation of either the CheRiff or ChrimsonR. A mixture of AAV^2/Rh10^ Syn-ChrimsonR-tdT 8.1 × 10^12^ vg/mL and AAV^2/Rh10^ CMV-GFP 8.1 × 10^12^ vg/mL was used for testing the co-expression rate of two independently expressing viruses. Viruses were expressed for 4 days. Imaris 10.1.0 was used to analyze the co-expression rate of the two viruses (Fig. 2a, b). Cre and DAPI channels were turned on and about 25 nuclei with both cre and DAPI staining were randomly selected from each image. Then the mRuby channel was turned on and the percentage of Cre+ mRuby-spots was determined. Next, the selection was started from mRuby and DAPI channels, and the percentage of mRuby+ Cre-spots was determined.

#### Electrophysiology

The electrophysiology setup (Supp. Fig. 1) was based on an Olympus BX61W1 epifluorescence microscope, Axopatch 200B amplifier (Axon) and a multi-wavelength fiber-coupled light source (Mightex, WFC). We used a 40x/1.0 Plan-Apochromat objective (Zeiss) which illuminates a field of 557 μm diameter. Optical stimulation experiments were performed 7-10 days after transduction. Whole-cell patch clamp recordings with oSTDP induction were performed in serum-free medium (in mM: 99 % MEM (Sigma; M7278), 13 D-glucose, 109 NaCl, 2 MgSO_4_, 1.44 CaCl_2_, 1 L-glutamine (Gibco 25030), 0.006 ascorbic acid (Fluka 11140), 0.01 mg/ml insulin (Sigma I6634). pH 7.28, 310-318 mOsm/kg) to mimic in-incubator conditions. The intracellular solution for patch-clamping and electroporation contained (in mW): 135 K-gluconate, 4 MgCl_2_, 4 Na2-ATP, 0.4 Na-GTP, 10 Na2-phosphocreatine, 3 ascorbate, and 10 HEPES, pH∼7. 2 (solution was sterile filtered, aliquoted and stored at - 20℃). CA1 pyramidal cells were voltage-clamped at −70 mV (liquid junction potential corrected), EPSCs were evoked by 594 nm laser pulses (2 ms, 0.05 Hz) directed through the condenser at CA3. After a short baseline period (< 5 min after break-in), oSTDP was induced. During plasticity induction (one 594 nm 2 ms light pulse (8 mW/mm^2^) paired with three 405 nm 2 ms light pulses (1.2 mW/mm^2^, 50 Hz), repeated 300 times at 5 Hz), the amplifier was switched to current clamp mode to allow spiking of the postsynaptic neuron. Next, the recording was switched back to voltage-clamp and further EPSCs were recorded (2 ms light pulses at 594 nm, 0.05 Hz) for 30 min. Series resistance and holding current were constantly monitored for quality control. ChrimsonR-induced spiking was assessed in cell-attached mode. EPSC amplitudes, slopes, and number of spikes were quantified using custom MATLAB scripts (R2016b).

#### Optogenetic induction of spike-timing-dependent plasticity

In-incubator oSTDP was performed according to published precedures^4^. Briefly, 1-2 days before stimulation, the last medium changed was performed and the slice culture was centered in a 35 mm culture dish. A stimulation tower with two LEDs was placed in the incubator. Two-color optical stimulation (identical to oSTDP induction during whole-cell recording) was controlled by a Master-8 stimulator (AMPI). Three days after in-incubator oSTDP, whole-cell recordings were performed in ACSF (in mM: 119 NaCl, 11 D-glucose, 2.5 KCl, 1 NaH_2_PO_4_, 4 MgCl_2_, 4 CaCl_2_, pH = 7.4, 305-315 mOsm/kg) aerated with Carbogen (95% O_2_, 5% CO_2_) at 31°C. CA1 pyramidal cells were voltage-clamped at −70 mV (liquid junction potential corrected) to record EPSCs triggered by 2 ms light pulses (594 nm) at 0.05 Hz.

#### Immunohistochemistry

Slice cultures were fixed in 4% PFA for 30 min, washed 3 times for 10 min in PBS and incubated for 1 h in blocking medium (PBS supplemented with 10 % donor goat serum, 0.3 % Triton X-100, 0.2 % BSA). For images in Fig.2, anti-Cre mouse primary antibodies (Millipore MAB3120) were diluted 1:500, and anti-mouse Alexa Fluor 647 (goat) secondary antibodies (Invitrogen #A21236) were diluted 1:1000. After fixation and PBS washes, slices were incubated with primary antibody mix overnight at 4°C. The following day after 3 washes in PBS, slices were incubated with secondary antibody mix for 2 hours, followed by washing, DAPI staining and mounting. For images in Fig. 3a (bottom), anti-RFP (for ChrimsonR-tdTomato) and anti-GFP (for CheRiff-eGFP) conjugated camelid nanobodies (NanoTag) were diluted 1:250 in carrier solution (phosphate buffered saline (PBS) supplemented with 1% donor goat serum, 0.3% Triton X-100, 0.2% bovine serum albumin (BSA) to label boutons and spines (Fig. 2a). After overnight incubation with the antibodies at 4°C, slices were washed 2 times in PBS, stained with DAPI (1:10000) and mounted in Mowiol, prepared according to manufacturer’s protocol with anti-fading reagent DABCO.

#### Confocal and Stimulated Emission Depletion (STED) microscopy

Gated-2D-STED images (Fig. 2a) were acquired using an Abberior STED Expert line microscope with a 60x/1.4 P-Apo objective (Nikon). Fluorophores were excited with 561 nm and 640 nm laser lines. Fluorescence was collected by avalanche photodetectors through emission filters 615/20 (red channel) and 685/70 (far-red channel). Temporal gating was obtained with a Becker and Hickl SPC150 TCSPC board (8 ns window). Both channels (red, far-red) were acquired in confocal mode. In addition, the far-red channel was resolved with 2D-STED by use of a 775 nm depletion laser. Acquisitions were obtained sequentially with a 20 nm × 20 nm pixel size. Confocal images were the result of 6 accumulations, STED images of 18 accumulations acquired in line-mode. Confocal and STED images were merged in Fiji/ImageJ, the lookup table was linearly adjusted for each channel.

#### Sample preparation for electron microscopy

Each slice culture was fixed with 2.5% GA + 2% PFA at 37°C for 1 min, transferred to ice-cold 2.5% GA + 2% PFA for 1 h followed by 3 × 10 min cacodylate buffer wash (all wash steps below were performed with cacodylate buffer on ice. Slices can be kept in cacodylate buffer at 4°C for up to 3 weeks before DAB staining). Next, the slice was incubated with 20 mM glycine for 15 min, followed by 3 × 10 min wash. Endogenous peroxidase was blocked by 0.3% H_2_O_2_ incubation for 15 min, followed by 3 × 10 min wash. A metal-enhanced DAB substrate kit (Thermo Scientific) was used for DAB staining. Slice cultures were incubated with 1 x DAB reagent for 1 h to ensure full penetration before stabilized hydrogen peroxide solution (1:100) was added to initialize the DAB reaction. After 30-120 min of staining (depending on the expression level), slice cultures were 3 times washed and incubated with 3% GA overnight. The following day, the slice was washed again 3 times. Stained sections were micro-dissected to the CA1 or CA3 region and kept in cacodylate buffer at 4°C for up to 3 weeks. OsO_4_ staining was then performed. The sections were washed in cacodylate buffer 2 x 5 min, followed by 20 min of 1% OsO_4_ incubation on ice. Then, the sections were transferred to cacodylate buffer for washing for about an hour, with buffer replacement every 5 min.

To prepare for resin embedding, dissected specimens were dehydrated in an ethanol series (30%, 50%, 70%, 80%, 90%, 100% x 2, 15 min each), then moved to 100% propylene oxide (2 times, 10 min each). Next, specimens were pre-embedded in a 1:1 mixture of EPON 812 resin (Glycidyl ether hardener MNA 16.1 g, Glycidyl ether hardener DBA 8.025 g, Glycidyl ether accelerator DMP-30 0.5 g and Glycidyl ether 100 25.85 g) and Propylene oxide for 90 min, then transferred to a 2:1 mixture of EPON 812 resin and Propylene oxide for 2 h, then transferred to pure EPON resin and incubated overnight. The following day, embedded specimens were placed on a resin block and cured in an oven at 58°C for at least 48 h.

DAB staining results (Fig. 1, step 5 and 6) were imaged on a Zeiss Axiophot with 10x/0.25 Achroplan objective and DFK37AUX250 camera (The Imaging Source), the micro-dissection was based on the DAB signal. After OsO_4_ staining and embedding, semi-thin sections (∼500 nm) were collected for further trimming (Fig 2, step 7): only the region containing the expected structure was kept for the collection of serial sections (e.g. *stratum radiatum* below a labeled CA1 soma, Fig. 4).

A diamond knife was used for manual serial sectioning. Ultra-thin sections (50 nm) were collected on Coffer slot grids (0.4 x 2 mm). For achieving higher contrast, post-staining contrasting was performed. Each grid was placed on a drop of 100 µl uranyl acetate solution for 15 min, then washed in water, air dried in a grid box and moved to a drop of lead citrate for 1 min. After another washing and drying step, samples were ready for imaging. All contrasting steps were performed in a closed petri dish.

#### Transmission electron microscopy and ultrastructural analysis

TEM images were acquired on a JEM-2100Plus electron microscope (JEOL) at 200 kV acceleration voltage with a JEOL recorder and CCD camera system (EMSIS), or on a TEM 900 electron microscope (Zeiss) at 80 kV acceleration voltage with a 4K digital camera (Tröndle, Moorenweis, Germany). Magnification was verified in different samples from 2,000 x to 20,000 x. Panoramic overview images were obtained by tiling TEM images obtained at lower magnification (7,000 x), with 64 panoramas covering almost an entire ultrathin section (14,400 μm^2^). Tiles containing labeled dendrites were identified by eye and high-resolution images (20,000 x) of those regions were acquired. Synapses were identified by searching for postsynaptic densities opposed by active zones and were manually classified as unlabeled, pre-only, post-only, or double-labeled. Labeled presynaptic vesicles were distinguished from naturally occurring dense-core vesicles by their smaller size and characteristic gray value. If only a single labeled vesicle was identified in a bouton section, we checked consecutive sections or neighboring boutons of the same axon for additional labeled vesicles for confirmation. Identified synapses were morphometrically analyzed with ImageJ, the curvature of the axon-spine interface (ASI) was measured using the Kappa plug-in^40^.

### Statistical analysis

Statistical tests and sample sizes are provided in the figure legends. Results with p < 0.05 were considered statistically significant. Statistical tests were performed and data were plotted using GraphPad Prism 10.

## Supplemental information

Document S1: Figures S1–S6.

