## Supplemental Figures 1 - 6 for "Ultrastructural analysis of synapses after induction of spike-timing-dependent plasticity"

### Supplemental Information

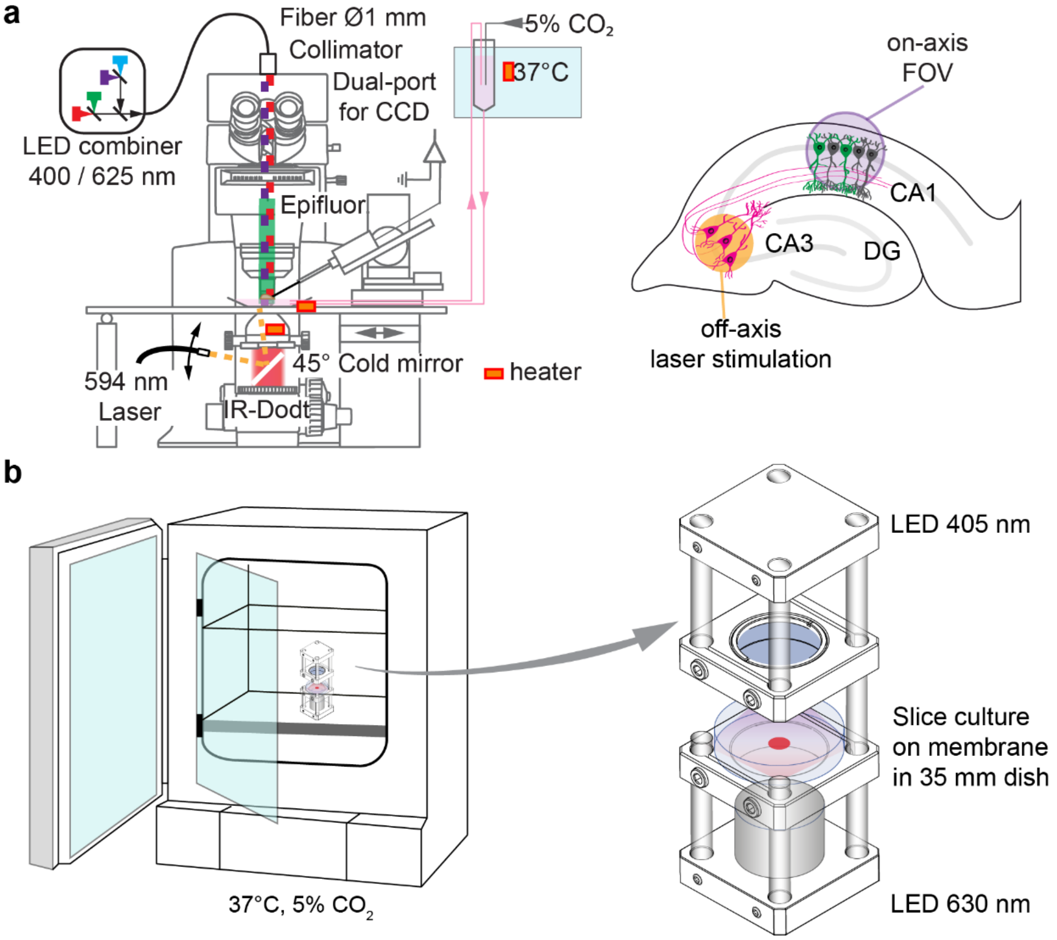


**Figure S1: Induction of STDP on the microscope stage and in the incubator.**

(**a**) For electrophysiology experiments, a microscope with a fiber-coupled LED combiner and a fiber-coupled yellow laser (for off-center illumination through the condenser) was used. (**b**) In-incubator stimulation. An LED tower was placed inside the incubator, delivering computer-controlled light pulses to the top (405 nm) and bottom (630 nm) of a 35 mm culture dish containing the membrane insert. Images adapted from: Anisimova et al. (2022) Cerebral Cortex 33, 23–34.

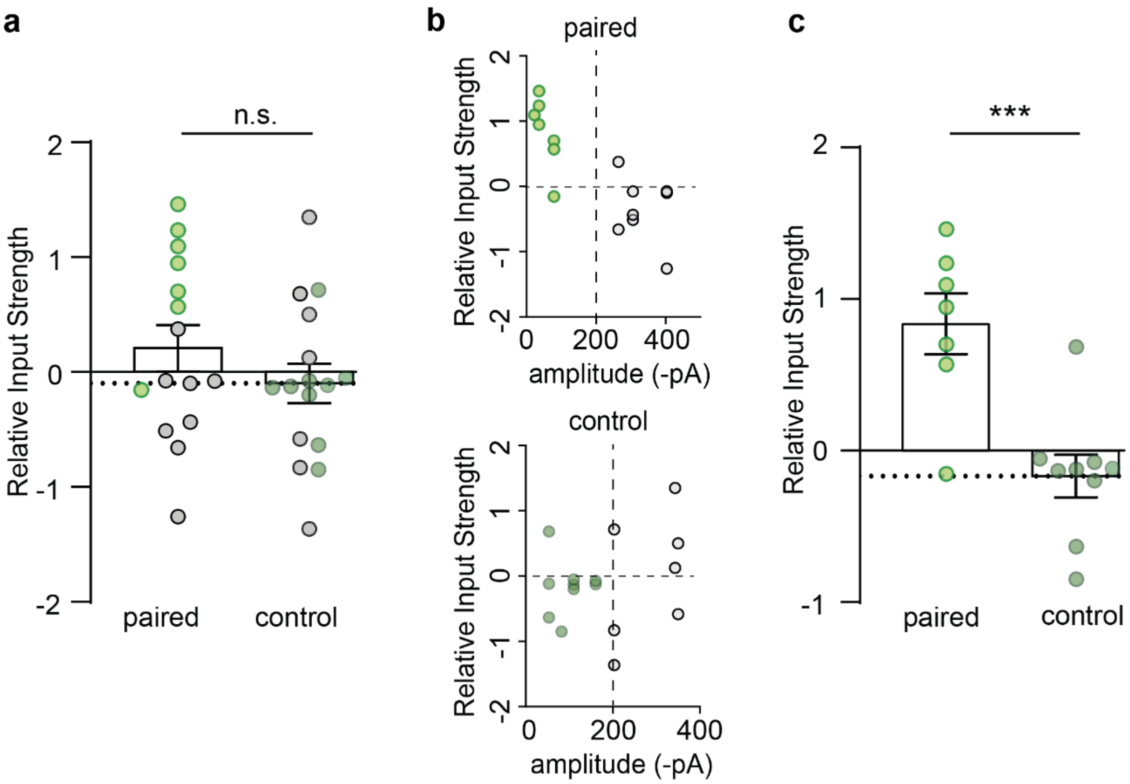


**Figure S2, related to Figure 3: Activating a large number of presynaptic neurons prevents selective potentiation of paired synapses.**

(**a**) Full dataset. Each dot indicates the EPSC amplitude of a CheRiff-expressing CA1 pyramidal cell relative to 2-3 non-transfected neighbors after causal pairing or without optical stimulation (control). No difference in input strength between the paired and the unpaired group relative to non-transfected neighbors. Green dots: average EPSC amplitude < 200 pA; black dots: average EPSC amplitude > 200 pA. n = 15, 16. Unpaired t-test, ns: not significant. (**b**) Replot of data in (a) to show the relationship of input strength versus the averaged NT amplitude in paired (upper) and unpaired (lower) groups. (**c**) All experiments with EPSC amplitude < 200 pA, indicating few ChrimsonR-expressing CA3 neurons. Significant input strengthening relative to non-transfected neighbors in the paired group only. n = 7, 9 neurons. Unpaired t-test, ***p < 0.001. Data plotted as mean ± SEM.

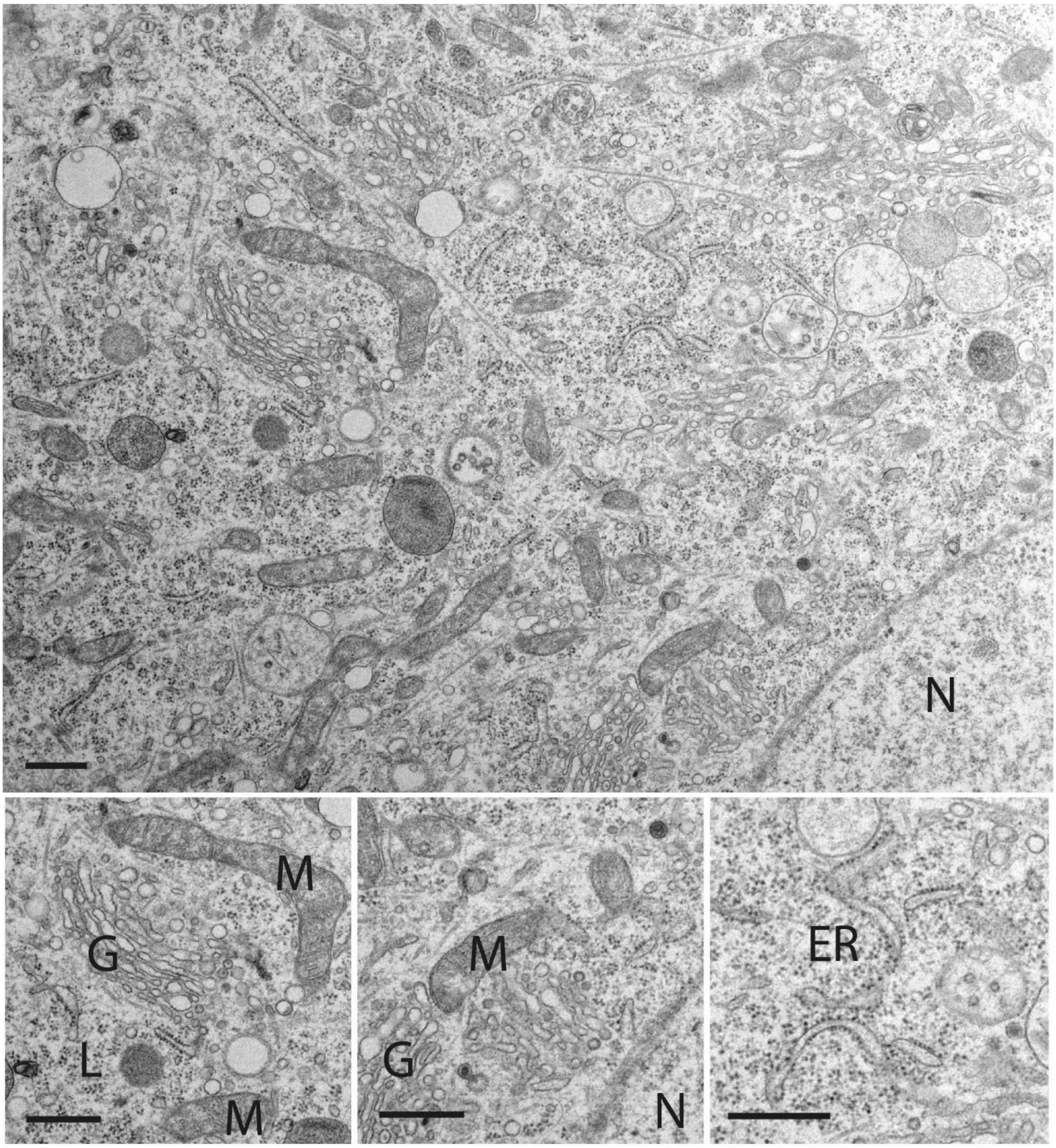


**Figure S3, related to Figure 4: Somatic ultrastructure of double-transduced neuron expressing SYP-HRP and ChrimsonR.**

Neuronal cytoplasm at the cell body including part of the nucleus (N). Due to synaptic vesicle targeting of HRP, no cytoplasmic DAB staining is visible in this neuron (see Fig. 4). Organelle morphology is well preserved (G: Golgi apparatus, M: mitochondria, L: lysosomes, ER: endoplasmic reticulum, N: nucleus). Scale bars: 500 nm.

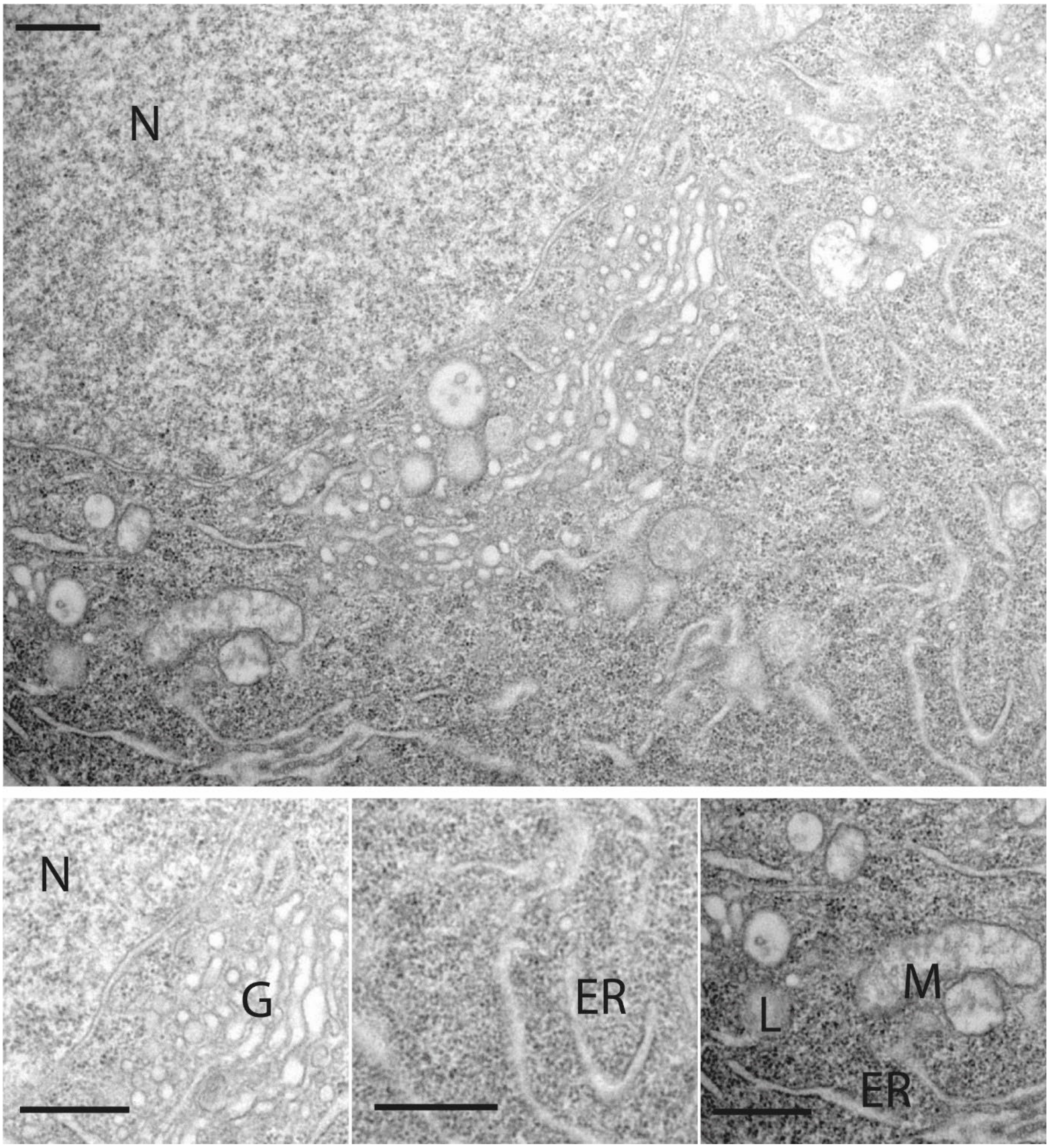


**Figure S4, related to Figure 4: Somatic ultrastructure of electroporated neuron expressing dAPEX2 and CheRiff.**

Neuronal cytoplasm at the cell body including part of the nucleus (N). Membrane contrast is relatively low due to dAPEX2 condensation in the cytoplasm (dark granules). Scale bar: 2 µm. Organelle morphology is well preserved (G: Golgi apparatus, M: mitochondria, L: lysosomes, ER: endoplasmic reticulum, N: nucleus). Scale bars: 500 nm.

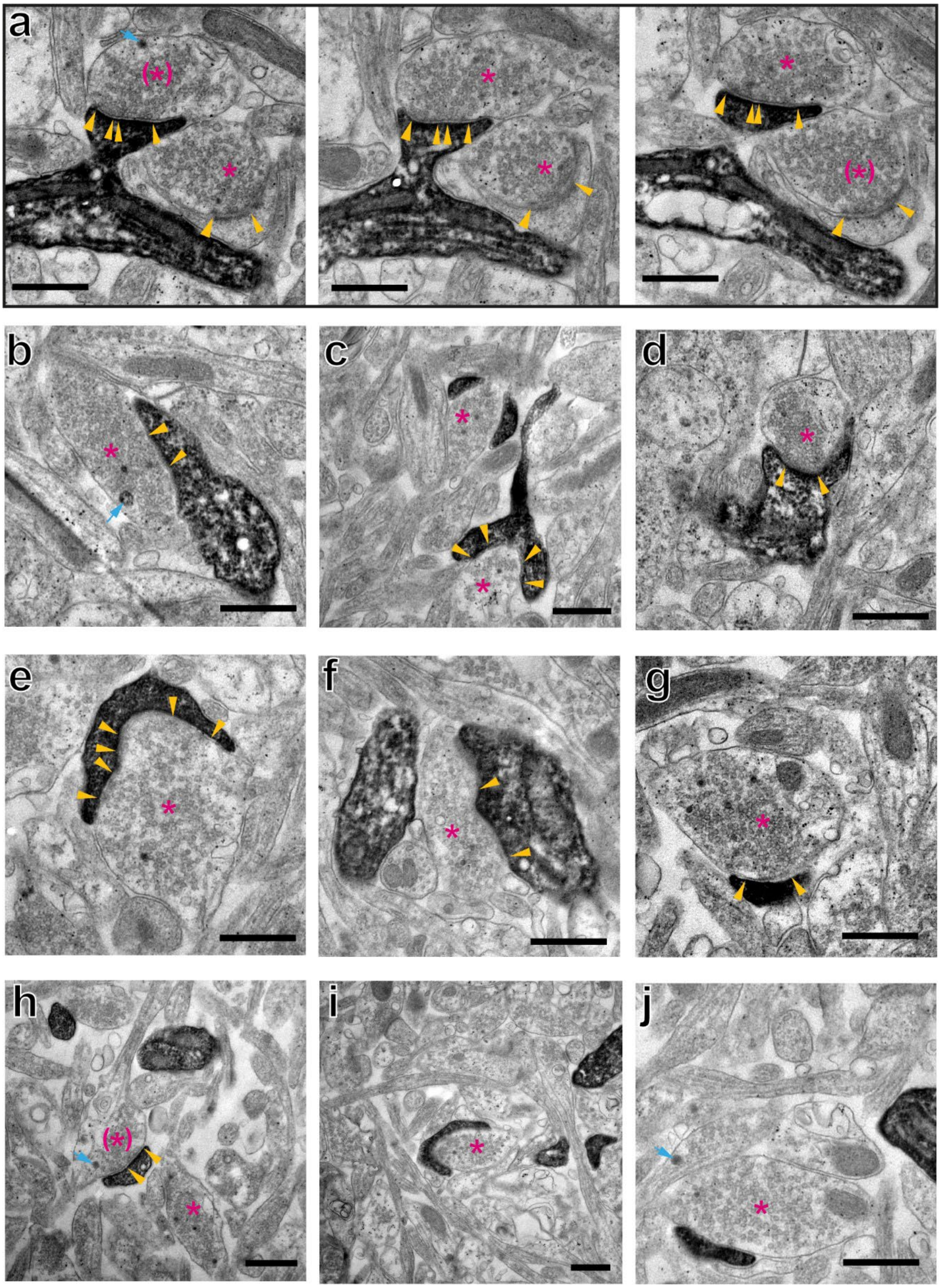


**Figure S5, related to Figure 4: Double-labeled contacts after causal stimulation**

(**a**) Two spine synapses identified in 3 consecutive sections. Yellow arrowheads mark edge of PSDs. Magenta asterisks: Stimulated presynaptic terminal containing one or more labeled vesicles. Magenta asterisks in brackets: Stimulated terminal identified by labeled vesicles in other sections. Cyan arrow: dense core vesicle, not indicative of labeling. (**b**) – (**h**) Further examples of double-labeled synapses, identified in single sections. (**i**), (**j**) Putative synapses, PSD not identified. All scale bars: 250 nm.

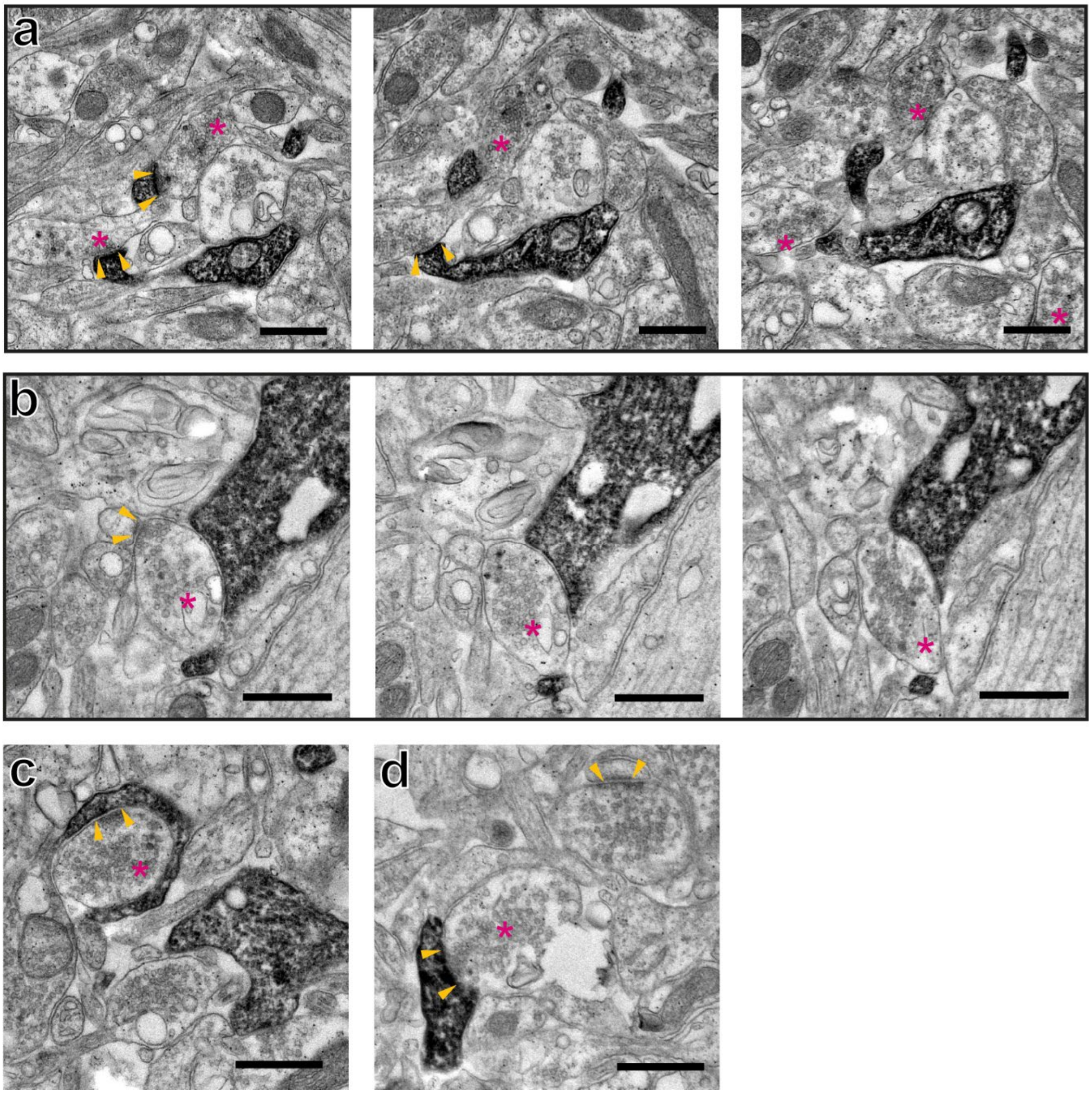


**Figure S6, related to Figure 4: Double-labeled contacts after anti-causal stimulation**

(**a**) Two double-labeled spine synapses identified in 3 consecutive sections. Magenta asterisks: Stimulated presynaptic terminal containing one or more labeled vesicles. Yellow arrowheads denote PSD. (**b**) Pre-only synapse (Yellow arrowheads) with contact to a labeled postsynaptic dendrite, identified in 3 consecutive sections. (**c**),(**d**) Further examples of double-labeled synapses. All scale bars: 250 nm.
